# Enzyme fatigue limits the detoxification of aflatoxin by *Rhodococcus* species

**DOI:** 10.1101/2021.08.21.457216

**Authors:** Marco Zaccaria, Natalie Sandlin, David Fu, Marek Domin, Babak Momeni

## Abstract

Bacterial detoxification of mycotoxins has the potential to offer a low-cost solution to ensure that feed and food commodities contaminated by fungal growth become safe to consume. Among bacteria, *Rhodococcus* species are of particular interest because they can be metabolically versatile, non-pathogenic, and environment-friendly. However, the native response of *Rhodococcus* environmental isolates appears inadequate for current detoxification needs. By analyzing the detoxification of aflatoxin by two *Rhodococcus* species: *R. pyridinivorans* and *R. erythropolis*, we examine important features of the dynamics that could guide future optimization of bacterial detoxification. Our results for *Rhodococcus* species suggest that detoxification happens through a regulated process of secreting extracellular enzymes. We show that enzyme fatigue in the presence of the toxin determines the lifetime of the enzyme and limits the overall detoxification performance of these species. Additionally, we show that the regulation of enzyme production can be both species- and environment-dependent. Overall, our quantitative approach reveals that enzyme fatigue is a major determinant of overall detoxification and needs to be accounted for in assessing the performance of detoxification by live cells or cell-free filtrates.

## Introduction

Aflatoxin is a secondary metabolite produced by species of the fungal genus *Aspergillus* (1), and it is widely regarded as the most dangerous among all mycotoxins—defined as toxic compounds of fungal origin. Aflatoxin contamination affects a wide range of food commodities, either directly (e.g. cereals, oilseeds, spices, and tree nuts) or indirectly (e.g. dairy and meat products of livestock exposed to the toxin) (2). Aflatoxins are among the most cancerous natural compounds, categorized as a Group 1 carcinogen by the International Agency for the Research on Cancer (IARC) (3). State-of-the-art decontamination relies on expensive and unsafe agrochemicals and costly production processes in the industrialized world, while developing countries still face enormous health risks (4). To protect consumers, large quantities of contaminated crops are disposed of across the globe every year, contributing to the large economic toll of mycotoxins: on the US economy only, an estimated $1.4 billion per year, and it is mostly attributable to aflatoxins (5). Aflatoxin intake primarily affects the liver (6) and is strongly associated with the insurgence of Hepatocellular Carcinoma (HCC) (7) by acting synergistically with Hepatitis B virus (8). Aflatoxin consumption is extremely frequent; in developing countries, about 5 billion people in 2006 consumed contaminated food (9): in societies where milk, cereals and nuts are central to everyday diet, exposure to high doses of aflatoxin is an unavoidable consequence (6).

Since aflatoxin is extremely stable in the environment, decontamination through physical and chemical means has been challenging in terms of cost, efficiency, and reliability (10). Bioremediation—the employment of live organisms to degrade the toxin—has emerged as a promising alternative (11–13) when specific requirements are satisfied: detoxification byproducts are safe for consumption and for the environment, microbes employed are harmless, and undesired side-effects are experimentally ruled out. Some species have been found capable of degrading aflatoxin (12, 14), including fungal species from the genera *Trichoderma* and *Rhizopus* (15) and bacterial species such as *Pseudomonas putida* (16), *Myxococcus fulvus* (17), *Enterococcus faecium* (18), and several *Rhodococcus* taxa (19, 20). The detoxification activity, in most cases, appears to be catabolic (21) and extracellular (22), although aflatoxin removal by adsorption is also possible, for example in lactic acid bacteria (23, 24).

At present, there are no identified wild type variants with adequate detoxification efficiency to be incorporated in the food production chain as a bioremediator. To achieve the required efficiency, optimization of detoxification performance (through artificial selection or other means) is likely required. However, it remains obscure where the hurdles are. Characterization assays typically measure the degree of toxin removal over a time interval (e.g. 24 or 48 hours). However, such assays do not reveal the true potential of each strain because they do not control for parameters such as enzyme density (a strain-dependent parameter) and enzyme kinetics. We propose to quantitatively infer such parameters to offer a systematic comparison of each strain’s detoxification performance and potential.

For our investigations, we use the bacterial genus *Rhodococcus*. This choice is made because there is an established record of aflatoxin detoxification by *Rhodococcus* species (20). Additionally, *Rhodococcus* species, often non-pathogenic and environment-friendly (25, 26), are amenable to genetic manipulations (27–29). Despite these advantages, progress towards real-world applications of these strains is hampered by lack of insights into the underlying mechanisms of detoxification. A thorough, mechanistic characterization of the detoxification process would provide solid basis to guide genetic engineering or artificial selection efforts for enhancing the detoxification performance. Our work focuses on two *Rhodococcus* species, *R. pyridinivorans* (*R-pyr*) and *R. erythropolis* (*R-ery*), which are among the most efficient species for degrading aflatoxin (20, 22, 30). We use the native fluorescence of aflatoxin G_2_ (AFG_2_) to quantitatively characterize how these strains detoxify aflatoxins in our investigations.

Our results reveal several important aspects. (1) Aflatoxin degrading enzymes of *Rhodococcus* species are under regulation and mostly produced during the mid-exponential growth phase. (2) Degrading enzymes lose efficacy during toxin degradation. (3) Using starch (compared to glucose) as the carbon source in the growth medium improves the degrading performance of *R-pyr* via an increase in enzyme production, but the same does not happen for *R-ery* strains. (4) The trade-off between enzyme concentration and lifetime during aflatoxin degradation limits the overall detoxification performance.

## Results

### 1. R. pyridinivorans and R. erythropolis detoxify AFG_2_ using extracellular enzymes

*Rhodococcus* species have been shown to hold ample biodegradation potential (25, 26). In particular, cell-free filtrates of *R-ery* were shown to degrade AFB_1_, pointing to extracellular enzymatic activity (22, 30, 31).

We tested the degradation of AFG_2_ by our *R-pyr* and *R-ery* filtrates after growing each strain to saturation (Fig 1). After 72 hours, filtrates of *R-pyr* cultures showed 71% ± 3% (s.d.) AFG_2_ degradation and, comparably, *R-ery* filtrates showed 80% ± 0.8% (s.d.). Examining the degradation profile of both species, it appears that the majority of degradation took place within the first 24 hours.

**Fig 1.**
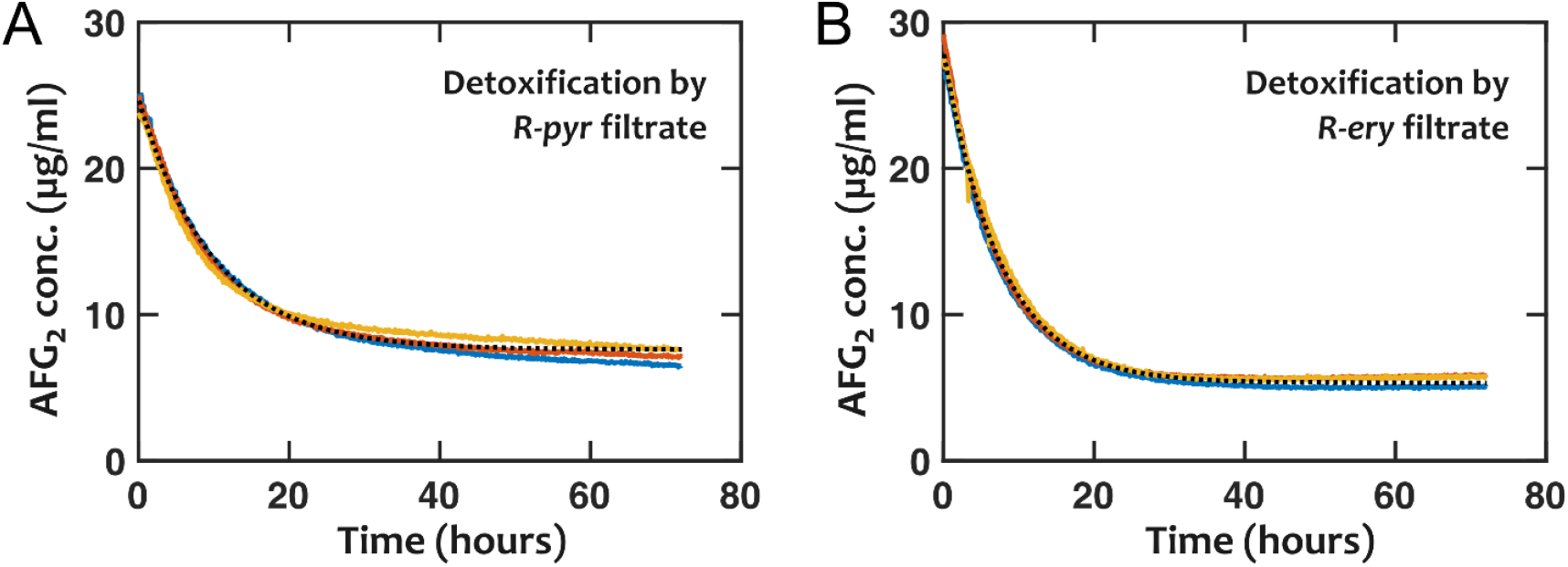
Detoxification of AFG_2_ by cell-free filtrates of *R-ery* and *R-pyr*. Each set of three replicates of *R-pyr* and *R-ery* filtrates (solid colors) shows consistent degradation profile, reaching AFG_2_ degradation of 71% ± 3% (s.d.) for *R-pyr* and 80% ± 0.8% (s.d.) for *R-ery*. Dotted lines show the inferred fit into the experimental data based on a Michaelis-Menten model with a fixed enzyme lifetime (Eq 1).

The slow-down in degradation by *R-pyr* and *R-ery* filtrates after 24 hours could be either because of a drop in the toxin concentration or the decay of the enzymatic activity. Since toxin concentration was still at a relatively high level—at 20-30% of its initial concentration after 24 hours—we ruled out the low toxin concentration as a major reason. Instead, when we fitted a model assuming a “fatigue” rate (corresponding to an enzyme lifetime) for the enzyme (Fig 1, dotted lines), the model was consistent with experimental data. In contrast, assuming no enzyme fatigue in the model led to results that were qualitatively inconsistent with our experimental observations (Figs S1 and S2).

For our model, we used a standard Michaelis-Menten description of enzyme kinetics to estimate the parameters associated with the extracellular enzymes of *R-pyr* and *R-ery* in their corresponding cell-free filtrates. Using the monitored toxin concentration over time, we obtained a progress curve that was the basis for our fitting of the model into the data. We used a minimal model with only two degrees of freedom: the initial enzyme concentration and the rate of enzyme fatigue.

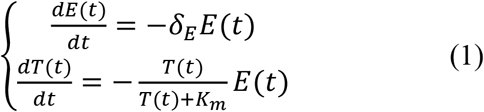

where *E*(*t*) is the enzyme concentration (in µU/ml), starting from *E*_0_ = *E*(*t* = 0) as the initial concentration of the enzyme in the filtrate, and *T*(*t*) is the concentration (in µg/ml). Two additional parameters determine the kinetics of the detoxification: *K*_*m*_ is the Michaelis-Menten coefficient and *δ*_*E*_ is the enzyme “fatigue” rate (i.e. enzyme decay rate in the presence of aflatoxin). To estimate *K*_*m*_, we monitored the detoxification of different initial concentrations of AFG_2_ (Fig S3) and estimated the *K*_*m*_ as the concentration at which the initial slope (0.5-1.5 hours, before enzyme fatigue could have a major impact) was half of its maximum. We later found that the exact value of *K*_*m*_did not play a critical role in our main findings, and our overall conclusions were robust against changes in estimated *K*_*m*_ values.

We first asked whether the enzyme itself would decay under the experimental conditions in the absence of the toxin. When we tested the degradation by filtrates stored for 0, 48, and 96 hours at 28°C (our experimental conditions), there was only a modest decrease in detoxification performance (Fig S4). We thus concluded that interaction with aflatoxin was responsible for the apparent limited lifetime. We speculate that the enzyme-toxin interaction may lead to this enzyme “fatigue”; for example, the substrate might not be released from the enzymatic pocket post detoxification, or reaction products may compromise subsequent enzymatic activity. To represent this effect, we define an effective enzyme lifetime as the inverse of the fatigue rate (*τ*_*f*_ = 1/*δ*_*E*_).

### 2. Compared to R-pyr, R-ery filtrates have higher initial enzyme activity

Even though the two filtrates demonstrate comparable overall aflatoxin detoxification performance (Fig 1), our analysis reveals underlying differences between the two producing strains. Compared to *R-pyr, R-ery* shows higher initial effective enzymatic activity (Fig 2A). This could be a result of higher enzyme concentration (possibly due to a larger enzyme production rate by *R-ery*), and/or the enzyme produced by *R-ery* may simply be a better AFG_2_ degrader. The higher initial activity of *R-ery* filtrate is partially offset by a shorter enzyme lifetime (Fig 2B), although the difference in lifetimes was not significant in our data. Overall, *R-ery* filtrates lose potency faster than *R-pyr* filtrates over time during detoxification. The combination of these two effects ultimately leads to a comparable overall detoxification efficiency in the examined 72-hour time interval.

**Fig 2.**
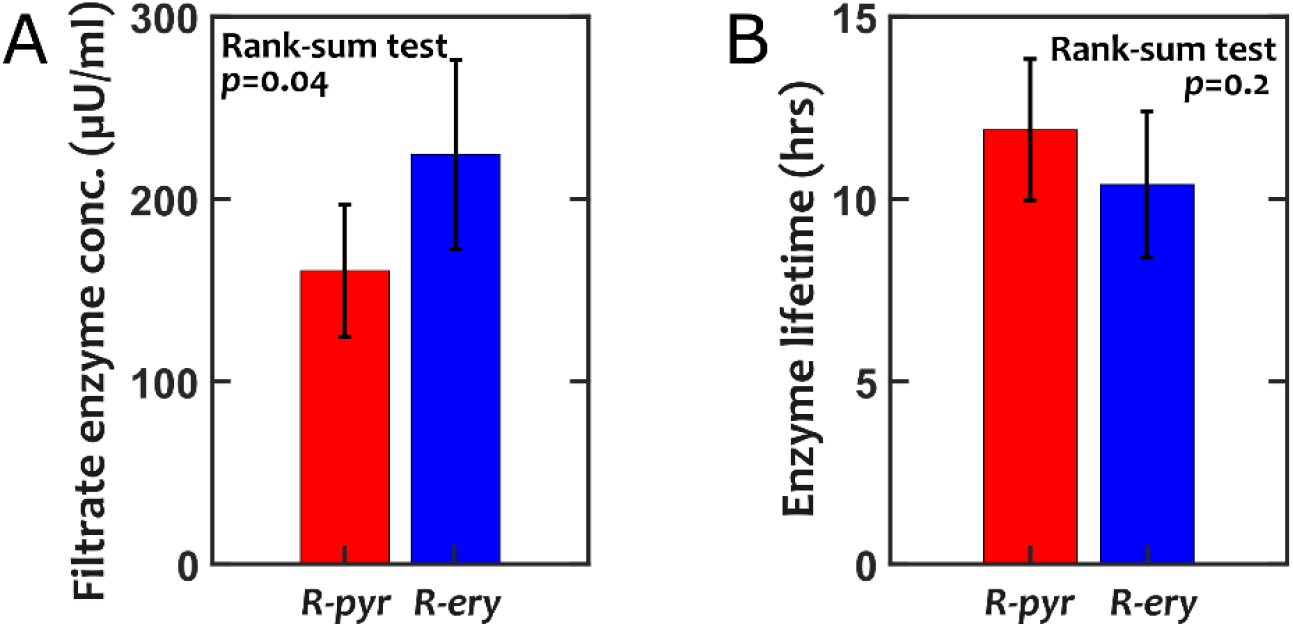
Filtrates of *R-pyr* and *R-ery* exhibit a trade-off between initial degradation efficiency and the rate of decay in enzymatic activity during the detoxification of AFG_2_. In our model, a constant fatigue rate was assumed for the enzyme throughout the detoxification process. Enzyme concentration is presented in μU/ml, with the definition that 1 U catalyzes 1 μmole of the substrate per minute (or 19.8 mg/hr of AFG_2_ with a molecular weight of 330 g/mole). Error-bars are s.d. obtained from three independent experiments with 2-3 replicates each.

The results in Fig 2 were obtained from three independent experiments. We observed that the variation was typically higher between experiments (run on independently isolated filtrates) rather than within experiments. This suggests that the detailed timing and experimental conditions for isolating the filtrate can have an impact on how the filtrate performs for detoxifying aflatoxin. Nonetheless, the overall trend between the two species remained the same: compared to *R-pyr* filtrates, *R-ery* filtrates initially showed stronger potency for degrading AFG_2_, but their performance also faded out faster.

### 3. Conventional definition of degradation efficiency is not an adequate measure for comparing different strains

Conventionally, the degradation efficiency of a toxin-removal process is defined as the fraction of the toxin degraded in a given amount of time (e.g. 60% in 24 hours). While this conveniently offers a single figure to describe the detoxification performance, we find that such a measure is sensitive to the details of the experimental setup. In Fig S5, we show that degradation efficiency is particularly sensitive to the initial toxin concentration in the test. This effect is especially exaggerated when the lifetime of the enzyme is short. As a result, we suggest that reporting the effective enzyme concentration and lifetime offers a more reliable representation of the detoxification performance.

### 4. The release of detoxifying enzymes by R-pyr and R-ery is regulated

To estimate the rate of the release of the enzyme, we use the following equations, with population density *S*(*t*)(in OD) and AFG_2_ concentration *T*(*t*) (in µg/ml) as observed quantities:

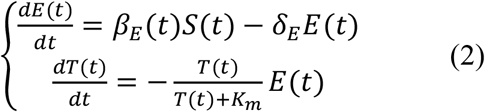

Here, *E*(*t*) is the effective concentration of the detoxifying enzyme over time (in µg/ml/hr) and *β*_*E*_(*t*) is the rate of enzyme release per cell. Rearranging Eq (2), we can calculate

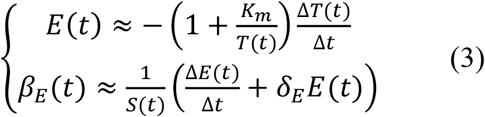

where Δ*t* is the sampling time step and Δ*T* and Δ*E* are changes in the AFG_2_ concentration and the effective enzyme concentration between time steps, respectively. From Eq (3), the temporal changes in *E*(*t*) and *β*_*E*_(*t*) can be inferred from the observable *S*(*t*) and *T*(*t*) dynamics. We find that, for both *R-pyr* and *R-ery*, the production of aflatoxin degrading enzymes is regulated (i.e., *β*_*E*_(*t*) is not a fixed number per cell during different stages of growth; Figs S1 and S2).

We ask whether the two species similarly regulated their enzyme release. To answer this question, we used Eq 3 to calculate enzyme release rate (per cell) for both *R-pyr* and *R-ery* (Figs 3 and 4, respectively). For both strains, the analysis shows that the production of the detoxifying enzymes is regulated, reaching a maximum rate in the mid-exponential growth phase, before tapering down subsequently. However, differences arise between the two strains in the early exponential phase, where the enzyme release rate appears has stable plateaus in the early and late exponential stages for *R-pyr* populations, whereas it increases towards a peak and decreases afterwards for *R-ery* (Figs 3D and 4D). Additionally, the per-OD rate of enzyme release appears to be higher for *R-ery*, even though this effect is countered by the overall lower final OD of *R-ery* cultures compared to *R-pyr* cultures in our experiments.

**Fig 3.**
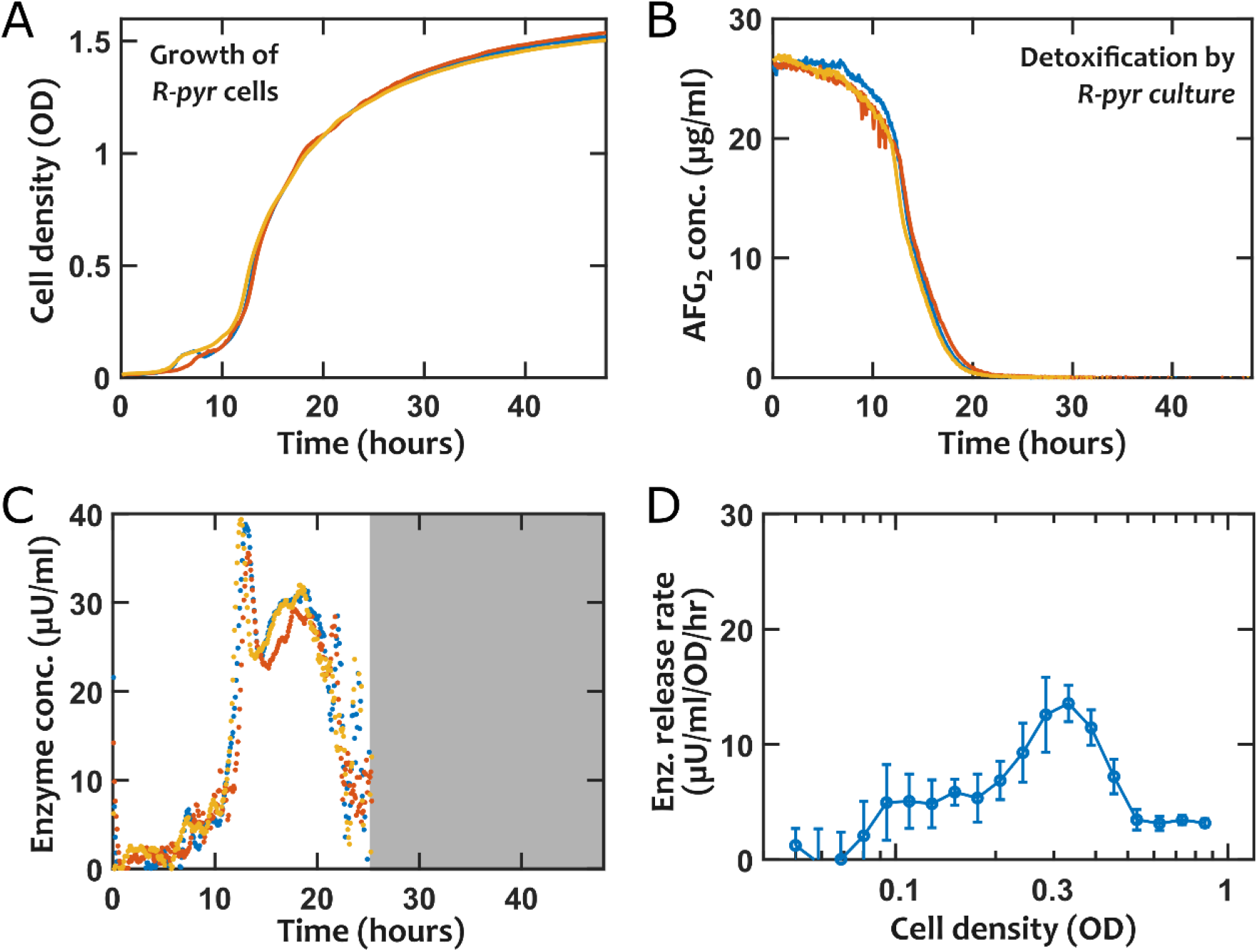
Probing the dynamics of AFG_2_ detoxification in live cultures of *R-pyr* suggests that the release of enzyme by cells depends on the growth stage of the culture. In (A) the total OD (at 600 nm wavelength) is used as a proxy for the cell density. In (B) the changes in AFG_2_ concentration is shown over time after converting the fluorescence reading to the corresponding concentration using the calibration curve. In (C) and (D) the inferred enzyme concentration and enzyme release rate are shown, respectively. The shaded region in (C) is excluded, because the toxin concetration was too low for reliably estimating the enzyme dynamics. In (D), data was binned into logarithmacially spaced groups based on OD, and the mean and s.d. of each bin is plotted.

**Fig 4.**
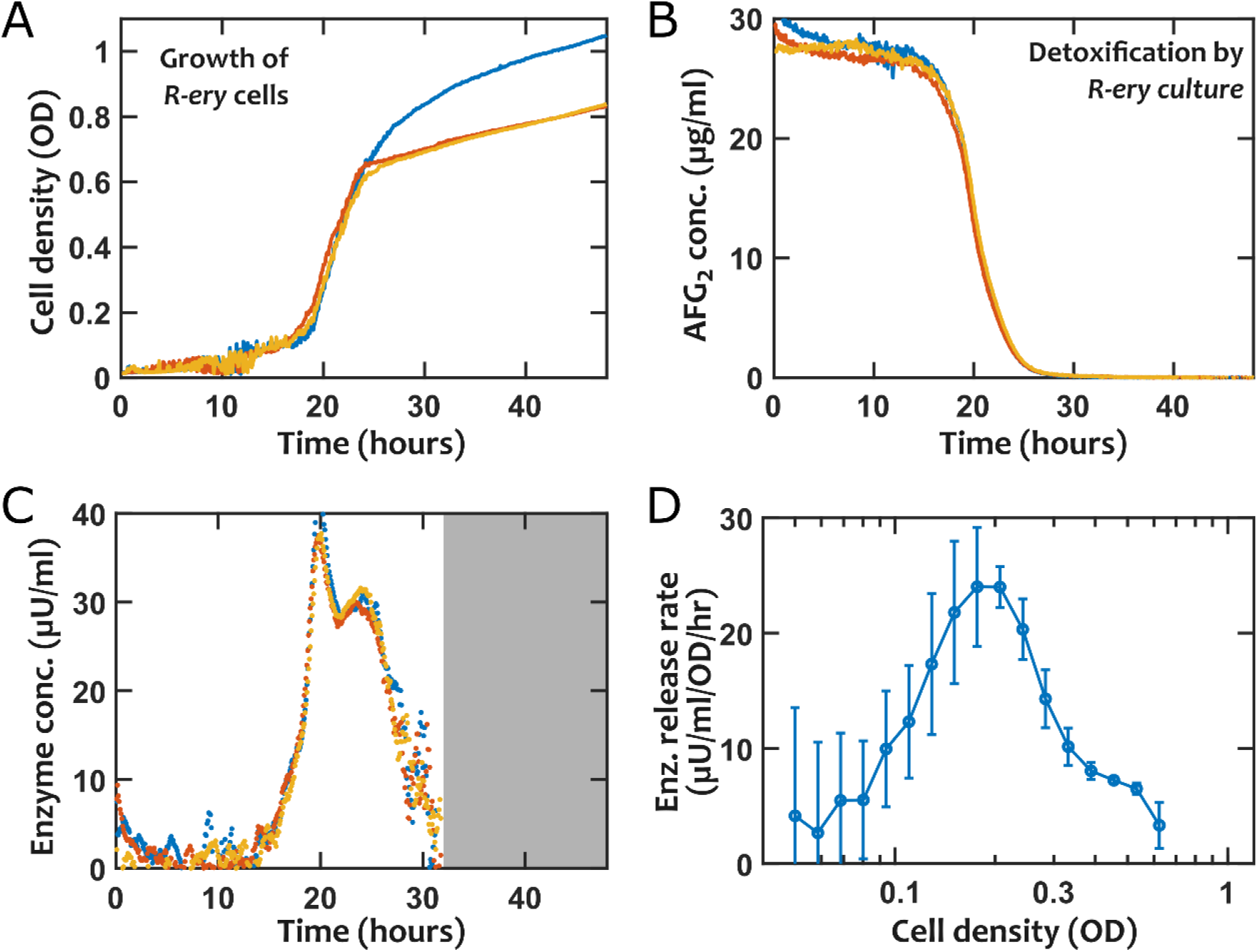
Probing the dynamics of AFG_2_ detoxification in live cultures of *R-ery* suggests that the release of enzyme by cells depends on the growth stage of the culture. In (A) the total OD (at 600 nm wavelength) is used as a proxy for the cell density. In (B) the changes in AFG_2_ concentration is shown over time after converting the fluorescence reading to the corresponding concentration using the calibration curve. In (C) and (D) the inferred enzyme concentration and enzyme release rate are shown respectively. The shaded region in (C) is excluded, because the toxin concetration was too low for reliably estimating the enzyme dynamics. In (D), data was binned into logarithmacially spaced groups based on OD, and the mean and s.d. of each bin is plotted.

We infer from our observations that: (1) the enzyme is likely involved in scavenging for resources in the environment, as it is upregulated when the cell runs low on readily accessible resources; and (2) the production/release of the enzyme is likely costly to the cell, as its production is regulated.

The observation that the rate of enzyme release depends on the growth stage has practical implications. For instance, to produce the detoxifying enzyme, a bioreactor could be maintained at the proper growth stage to maximize the enzyme release rate (Figs 3D and 4D).

### 5. The presence of starch as an alternative carbon source in the environment impacts detoxification by R-pyr *but not* R-ery

In search for potential environmental cues that could modulate the production of the detoxifying enzymes, we hypothesized that these enzymes may be involved in scavenging complex carbohydrates. This is motivated by the observation that enzyme production increased from early-to mid-exponential phase (Figs 3 and 4), when presumably the most accessible carbon sources are lowered. To test this, we replaced the glucose in our growth medium with the same carbon molarity of starch. Similar to Fig 2, we estimated the initial enzyme concentration and the enzyme decay rate for filtrates of *R-pyr* and *R-ery* obtained from starch cultures (Fig 5). Filtrates isolated from *R-pyr* cultures in starch medium showed an increase in the detoxifying enzyme concentration compared to the glucose-based medium; the same could not be said for *R-ery* (Fig 5A). Using Wilcoxon rank-sum nonparametric test, we found that the detoxifying enzyme concentration in *R-pyr* filtrates was significantly different between filtrates isolated from starch versus glucose medium (*p* = 0.001). In contrast, *R-ery* filtrates did not show a significant difference (starch versus glucose, *p* = 0.4). Unpaired two-sample t-tests corroborated with these rank-sum tests (with *p* = 9 × 10^−6^ and *p* = 0.6, respectively).

**Fig 5.**
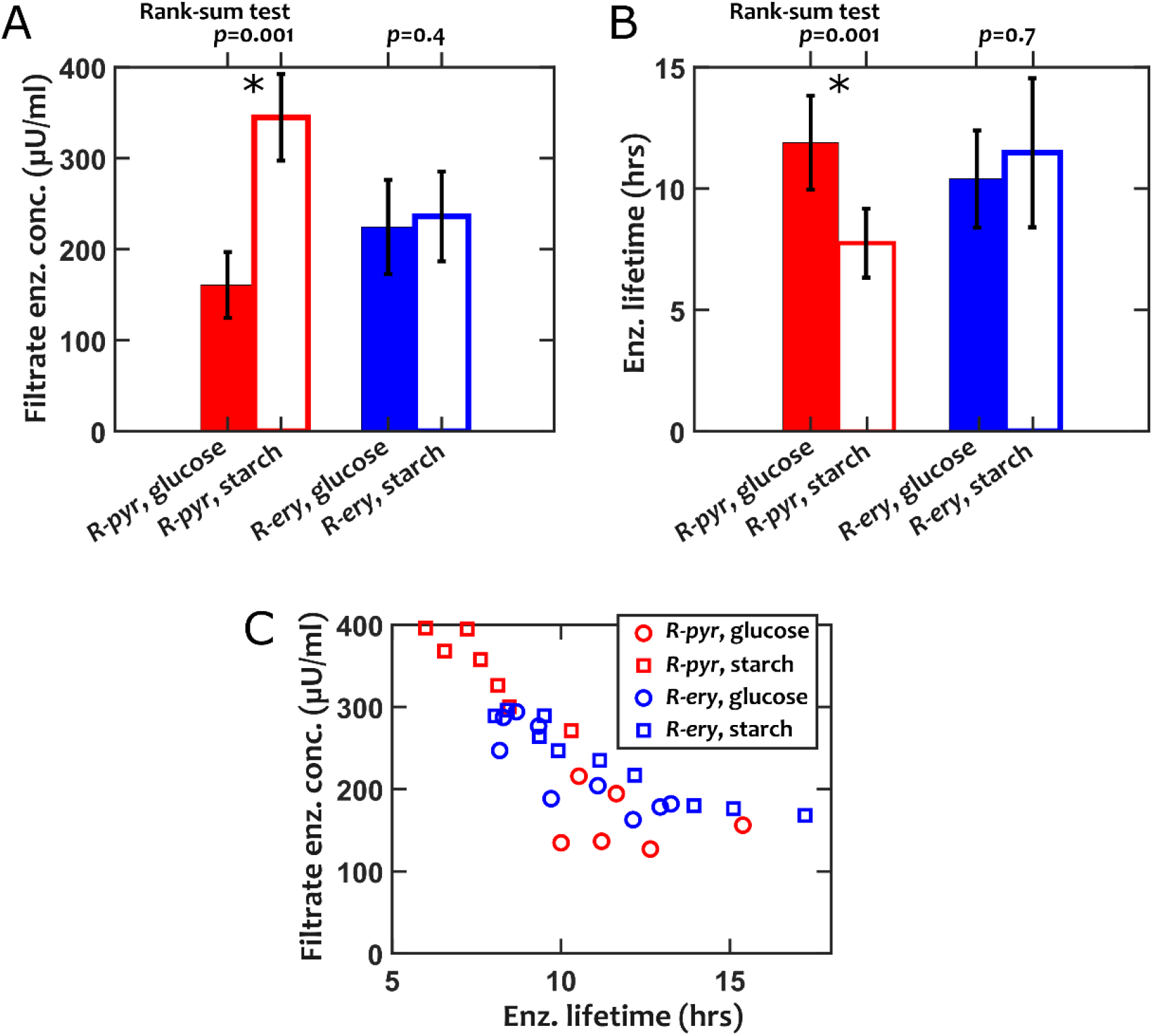
AFG_2_ detoxification is improved when using filtrates of *R-pyr*, but not *R-ery*, obtained from growth on starch (compared to glucose) as the primary carbon source. (A) Solid bars and open bars show the estimated concentration of the AFG_2_ degrading enzyme of filtrates taken from glucose and starch media, respectively. (B) Solid bars and open bars show the estimated enzyme lifetime of filtrates taken from glucose and starch media, respectively. (C) Filtrate enzyme concentration and lifetime of both *R-pyr*, and *R-ery* filtrates appear to show a consistent trade-off, with higher enzyme concentrations exhibiting a lower enzyme lifetime. We estimate this relationship as *τ* = 1/(4 × 10^−4^*E*_0_) with *τ* in hours and *E*_0_ in µU/ml (Fig S6). Data points are collected from three independent experiments. Error-bars are s.d. values and Wicoxon rank-sum test (a nonparametric comparison of medians) is used for statistical comparisons.

We speculate on three possible explanations: (1) Glucose as the preferred carbon source represses the production of relevant enzymes (a mechanism often observed in diauxie). (2) Starch induces the production of enzymes incidentally relevant to aflatoxin detoxification (similar to gal induction). (3) Starch supports more growth and in turn an increase in enzyme production. Additional investigations are needed to determine which mechanism is responsible for the change in AFG_2_-detoxifying enzyme concentration of *R-pyr* and how prevalent that mechanism might be among other aflatoxin-detoxifying species.

Since higher enzyme concentrations in our starch filtrates also exhibited lower enzyme lifetimes (Fig 5B), we asked if there was a trade-off. We examined all data-points (starch versus glucose filtrates and *R-pyr* versus *R-ery*), and the results showed a consistent trend (Fig 5C). Although not conclusive, Fig 5C suggests that the detoxification enzyme between *R-pyr* and *R-ery* is potentially similar, and that there is an intrinsic trade-off that a higher enzyme concentration lowers the detoxifying enzyme lifetime.

### 6. The limits of detoxification performance can be estimated from characterized parameters

We recognize that the environmental context (e.g. as observed in Fig 5) influences the observed detoxification performance. We also recognize that rarely the native strains found in nature will be directly employed for detoxification. Here we ask instead what is the expected performance of the strains we have characterized. For this, we operationally define “time to 95% detoxification” as a measure of the detoxification performance of the enzymes. This measure depends on the initial effective enzyme concentration, the initial toxin concentration, and the trade-off between enzyme concentration and enzyme lifetime. Since the trade-off for *R-pyr* and *R-ery* followed the same trend, we approximate this (Fig S6) as *τ*_*f*_ ≈ 1/(4 × 10^−4^*E*_0_) (in hours), where *τ*_*f*_ is the enzyme lifetime, and *E*_0_ is the initial enzyme concentration in the filtrate (in *μU*/*ml*).

Our results show that the trade-off in enzyme lifetime leads to a distinct limit in the detoxification performance (Fig 6A). Examining different initial enzyme concentrations in the filtrate (Fig 6B), we see that increasing the enzyme concentration only improves the performance at low aflatoxin concentrations and exhibits diminishing returns (because of the shorter enzyme lifetime) at higher concentrations. We repeated this analysis with parameters estimated for small *K*_*m*_ and observed similar trends (Fig S7).

**Fig 6.**
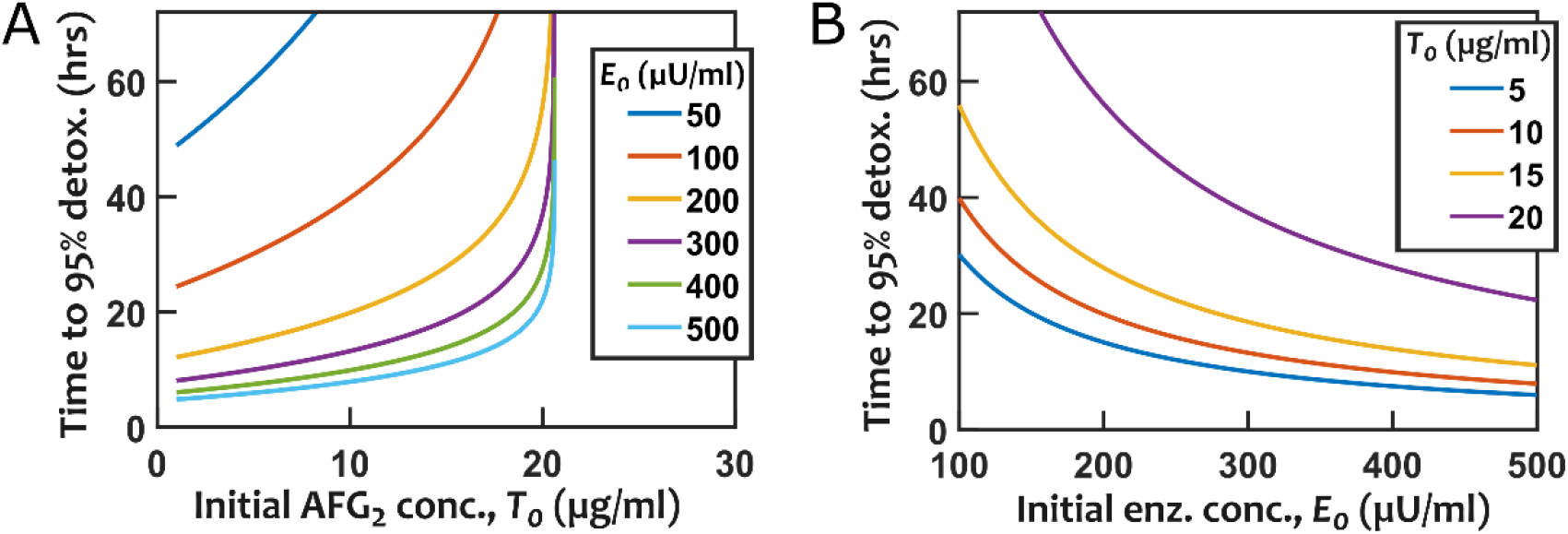
Simulated detoxification performance based on the time to detoxify 95% of the initial aflatoxin shows that enzyme fatigue limits the overall detoxification potential. The trade-off between enzyme concentration and enzyme lifetime limits the detoxification performance. (A) For each distinct enzyme concentration, detoxification time rapidly increases beyond a threshold aflatoxin concentration. (B) The detoxification of large aflatoxin concentrations exhibits diminishing returns at high enzyme concentrations. Eq (2) is used for these simulations, with the trade-off incorporated as a linear interpolation (Fig S6). *K*_*m*_ is assumed to be 10 µg/ml.

## Discussion

Bioremediation of aflatoxin contaminated food is an important area in food safety with global economic and health impacts (32, 33). Our work, in agreement with previous studies, illustrates that *R-pyr* and *R-ery* are capable of detoxifying AFG_2_ through extracellular enzymatic activity. Filtrate obtained from fully grown cultures showed detoxification potential of 70-80% in 72 hours (Fig 1), whereas live cultures were capable of completely removing AFG_2_ within 24 hours (Figs 3 and 4). We also find that the release of the enzyme is regulated, increasing and reaching a maximum in the mid-exponential phase, before decreasing again as cultures move to the stationary phase (Figs 3 and 4). We speculate that the enzyme is likely related to utilizing more complex carbon sources, and show an increase in the released enzyme by *R-pyr* when starch, rather than glucose, is the primary carbon source. Filtrates of both and *R-ery* also showed a consistent trade-off between the amount of the enzyme in filtrates and the enzyme lifetime. This trade-off ultimately limits the performance of enzymatic detoxification, especially for large aflatoxin concentrations. These insights inform future efforts towards improving the efficiency of aflatoxin biodegradation.

While bacterial species such as *R-ery* and *R-pyr* have the capability to degrade aflatoxin, it is unlikely that their enzymes are selected for optimal aflatoxin detoxification. This is because the inhibitory fitness impact of aflatoxin on bacteria is small (data not shown), and aflatoxin concentration is relatively low in the environment, making it unlikely that aflatoxin is used as a resource for growth. We thus propose that there is room for improved bioremediation performance if bacteria (or their enzymes) are selected or engineered for aflatoxin detoxification.

We did not observe a major difference between *R-ery* and *R-pyr* in terms of their AFG_2_ detoxification performance. The strains we tested also showed a consistent trade-off (Fig 5C), suggesting that detoxification of aflatoxin by these strains likely takes place through a similar mechanism. It would be interesting to perform detailed analysis of detoxification performance of additional strains and species in the future to see whether or not there are different categories of enzymes with distinctively different trade-offs.

Since the release of aflatoxin degrading enzymes in *R-pyr* and *R-ery* appears to be regulated, we also speculated that the presence of aflatoxin itself may contribute to the regulation. As a first estimate, we compared the concentration of the enzyme in the filtrates obtained without AFG_2_ exposure (Fig 2) with the final concentration inferred from live cultures exposed to AFG_2_. We calculated the latter by integrating the enzyme release rates over time (data from Figs 3 and 4; results in Fig S8). If AFG_2_ considerably modulated the release of the detoxifying enzyme, the total enzyme release until reaching the stationary phase should be different from what we observe in filtrates isolated from live cultures not exposed to AFG_2_. The enzyme concentrations estimated in *R-pyr* and *R-ery* filtrates without AFG_2_ exposure (Fig 2) were 160±36 µU/ml and 220±52 µU/ml, respectively. In comparison, the values inferred from live cultures (even after assuming an enzyme decay of 0.008/hr, estimated from Fig S4) were 320±15 µU/ml and 340±11 µU/ml, respectively (Fig S8). Inferred values from live cultures in the presence of AFG_2_ thus showed higher enzyme concentration, suggesting the possibility that the presence of AFG_2_ leads to a modest increase (< 2-fold) in the enzyme release.

Since the release of the detoxifying enzyme by *R-ery* and *R-pyr* appears to be regulated, we propose that this knowledge can be used to optimize the harvesting of the filtrates; keeping the growth stage in the maximal enzyme release range in a chemostat can lead to more efficient enzyme production, for instance. Additionally, the clear impact of regulation suggests that manipulating such regulation is a promising avenue for strain engineering in the future.

Lastly, progress in making bioremediation more efficient depends on better understanding of the underlying mechanisms. In particular, we emphasize that identifying, isolating, and characterizing the detoxifying enzyme should be a fruitful next step. These will also enable more directed future strain engineering efforts: on one hand, to uncover the important machinery that regulates the production and/or secretion of the detoxifying enzymes, and on the other hand, to allow potential transfer of the detoxifying enzymes to tractable organisms and/or to generally regarded as safe (GRAS) organisms for practical deployment.

## Materials and Methods

### Cultures and growth conditions

*Rhodococcus erythropolis* (DSM 43066) and *Rhodococcus pyridinivorans* (DSM 44555) were grown respectively in glucose-yeast-malt (GYM) and tryptic soy broth (TSB) at 28° C with continuous shaking (240 rpm) for 24 hrs. Experiments were led with both strains cultured in Basal Z medium: KH_2_PO_4_ (1.5 g/L), K_2_HPO_4_ × 3H_2_O (3.8 g/L), (NH_4_)_2_SO_4_ (1.3 g/L), sodium citrate dihydrate (3.0g/L), FeSO_4_ (1.1 mg/L), glucose (4.0 g/L), 100x vitamin solution (1 mL), 1000x trace elements solution (1 mL), 1 M MgCl_2_ (5 mL), 1 M CaCl_2_ (1 mL), and 100x amino acid stock (10 mL). Additionally, a modified Basal Z, Basal starch medium, was used replacing glucose (w/v) with starch. AFG_2_ (Cayman Chemical) was dissolved in LC-MS grade methanol to the final concentration of 1 mg/mL.

### AFG_2_ degradation assay

Cells or culture filtrates were aliquoted into in black glass-bottom 96-well plates (Nunc™ #165305 96-Well Optical Bottom). Aflatoxin was added according to desired final concentration per well. Final volume per well was 150 µl. Standard controls of toxin alone (AFG_2_ in fresh medium) and no toxin (cells or filtrate alone) were used. A BioTek Synergy Mx multi-mode microplate reader was used to monitor optical density of cells at 600 nm and fluorescence of aflatoxin at an excitation of 380 nm and emission of 440 nm with a gain of 50. Reads were taken at 5 min intervals over 72 hrs (unless otherwise noted). Cultures usually started at an initial OD of 0.01 and were continuously shaking between reads. Typically, 3-6 replicates were used per condition. Sterile water was placed at the peripheral wells of the plate to contain evaporation.

### Culture filtrates degradation assay

Basal Z or Basal starch liquid media (3 mL) were inoculated by frozen stocks of *R. erythropolis* and *R. pyridinivorans* and incubated at 28° C with shaking (240 rpm) for 24 hrs. OD of cultures was read before being filtered using 0.22 μm, PVDF syringe filters (Thomas Scientific). Filtrates were aliquoted into 5 mL sterile reservoirs and toxin was added for a final concentration of 15 ug/mL. Samples of filtrate with toxin were arrayed in a 96-well plate and underwent degradation assay as described above.

### Conversion of fluorescence read-out to aflatoxin concentration

For all cases, fluorescence is monitored using a fluorescence microplate reader (BioTek Synergy Mx) with 150 µl of sample in black glass-bottom 96-well plates (Nunc™ #165305 96-Well Optical Bottom). Fluorescence reads were obtained with excitation at 380 nm and emission at 440 nm both with 20 nm bandwidth. AFG_2_ was read using a gain of 50. Background fluorescence (no toxin control) is subtracted from the readings to remove fluorescence from sources other than the toxin. The readouts are also normalized to fluorescence data from no enzyme controls to remove the effect of fluorescence loss due to bleaching or other causes over time. The resulting values are then transformed to toxin concentration using a calibration data that contained values corresponding to known enzyme concentrations (34).

### Fitting Michaelis-Menten model into filtrate data

We used `lsqnonlin` routine in `Matlab®` to obtain the parameters that yielded the best fit into our experimental data. We used a simple transformation by choosing 0.1 *δE* (in 1/hr) and 1/50.5*E*_0_ (in µU/ml) as the free fitting parameters to make them comparable in magnitude and thus facilitate the optimization process.

## Code Availability

The following codes are used for fitting and analysis of data and are shared on GitHub for reproducibility at https://github.com/bmomeni/rhodococcus_aflatoxin_detox.

| Code | Application | Usage |
| --- | --- | --- |
| fit2K_Filtrate_AFG2.m | Fitting a model into filtrate degradation data | Figs 1 |
| func_EnzymaticDetox_Filtrate2K.m | Cost function for fitting a model into filtrate degradation data | Fig 1 |
| DetoxKinetics.m | Function that calculates the kinetics of detoxification by filtrates |  |
| DetoxKineticsCells.m | Function that calculates the kinetics of detoxification by live cultures |  |
| EstimateEnzReleaseRateK_Rery_AFG2.m | Infer enzyme concentration and release rate for live cultures of *R-ery* | Figs 3 and S8 |
| EstimateEnzReleaseRateK_Rpyr_AFG2.m | Infer enzyme concentration and release rate for live cultures of *R-pyr* | Figs 4 and S8 |
| PlotDegradationComparison2K.m | Comparing glucose versus starch filtrates | Figs 2, 5, and S6 |
| CompareTimeToDetox_InitialEnzConc.m | Calculate detoxification time by simulating the enzyme trade-off (over a range of filtrate enzyme concentrations) | Figs 6 and S7 |
| CompareTimeToDetox_InitialToxinConc.m | Calculate detoxification time by | Figs 6 and S7 |
|  | simulating the enzyme trade-off (over a range of initial aflatoxin concentrations) |  |
| Analysis_Rpyr_AFG2_Cells.m | Compare *R-pyr* experimental results with simulation results assuming no enzyme decay | Fig S1 |
| Analysis_Rery_AFG2_Cells.m | Compare *R-ery* experimental results with simulation results assuming no enzyme decay | Fig S2 |
| DetoxByFiltrate_AFG2_glucose_InitialToxConc.m | Examine the impact of initial toxin concentration on detoxification by *R-pyr* and *R-ery* filtrates to estimate $K_m$ | Fig S3 |
| StoredFiltrate_AFG2.m | Examine the detoxification performance of filtrates when stored in the absence of aflatoxin | Fig S4 |
| CompareDegEff_DiffInitialToxinConc.m | Simulate the dependency of degradation efficiency on the initial concentration of AFG<sub>2</sub> | Fig S5 |

## Acknowledgements

This work was supported by a start-up grant and an Ignite grant from Boston College and by a grant from the National Science Foundation (NSF-CBET, Award No. 2103545). NS is supported by a NIFA-AFRI predoctoral fellowship from USDA (Award No. 2021-67034-35108).

## Supplementary Information

**Fig S1.**
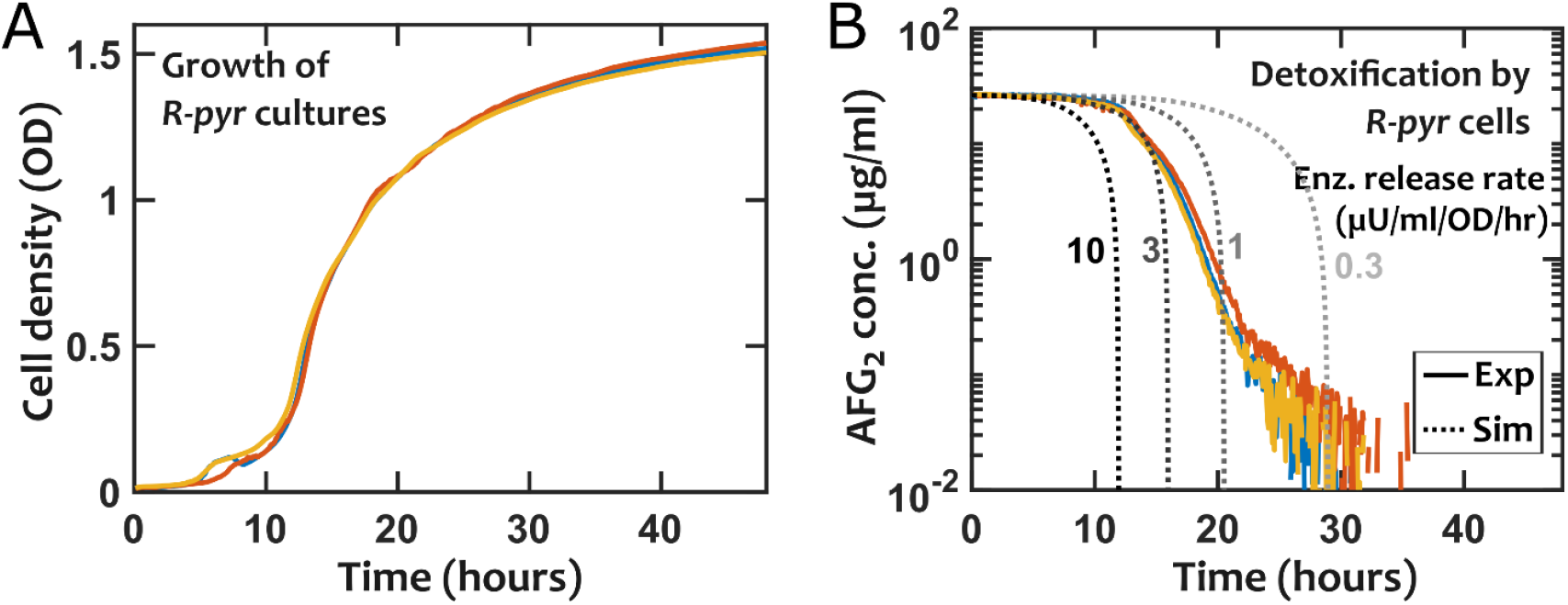
Detoxification of AFG_2_ by live cultures of *R-pyr* cannot be explained using a model that assumes cells release the enzyme at a fixed rate. (A) Measured OD is used as a proxy for cell density. (B) Using the measured OD and assuming a fixed enzyme release rate (shown on each graph in µU/ml/OD/hr) and assuming no enzyme decay.

**Fig S2.**
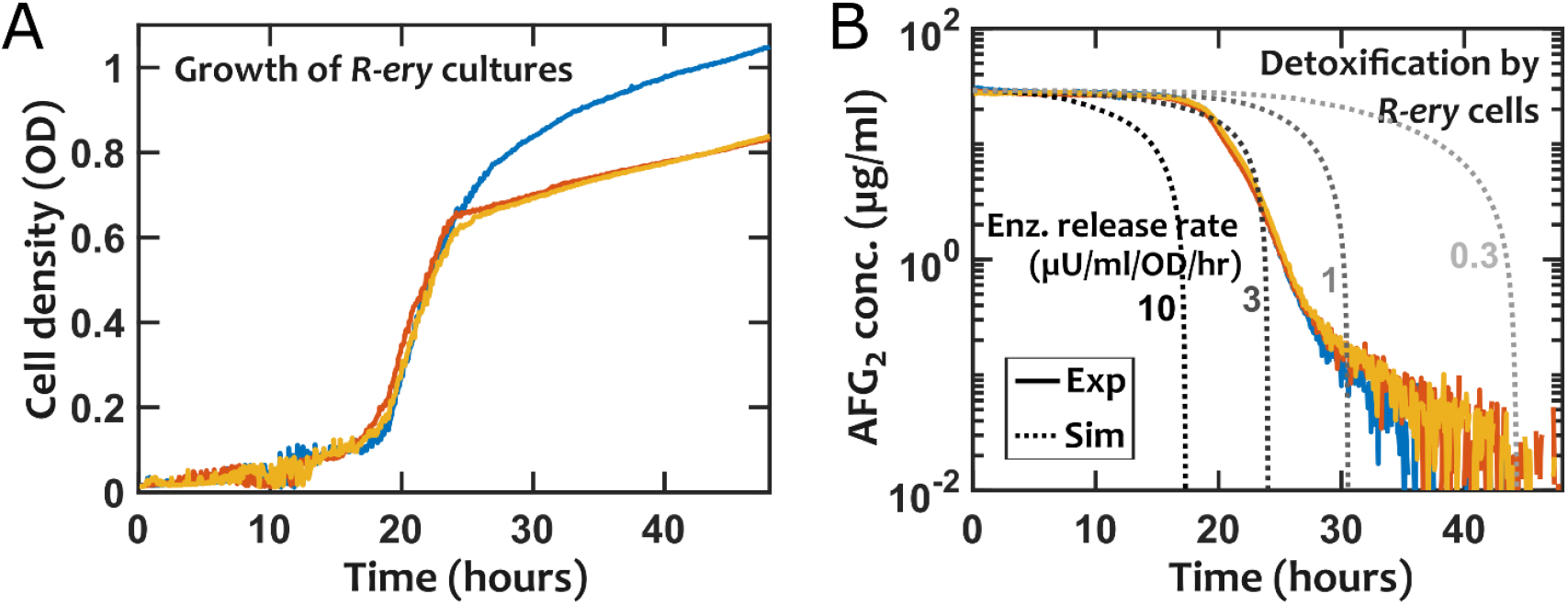
Detoxification of AFG_2_ by live cultures of *R-ery* cannot be explained using a model that assumes cells release the enzyme at a fixed rate. (A) Measured OD is used as a proxy for cell density. (B) Using the measured OD and assuming a fixed enzyme release rate (shown on each graph in µU/ml/OD/hr) and assuming no enzyme decay.

**Fig S3.**
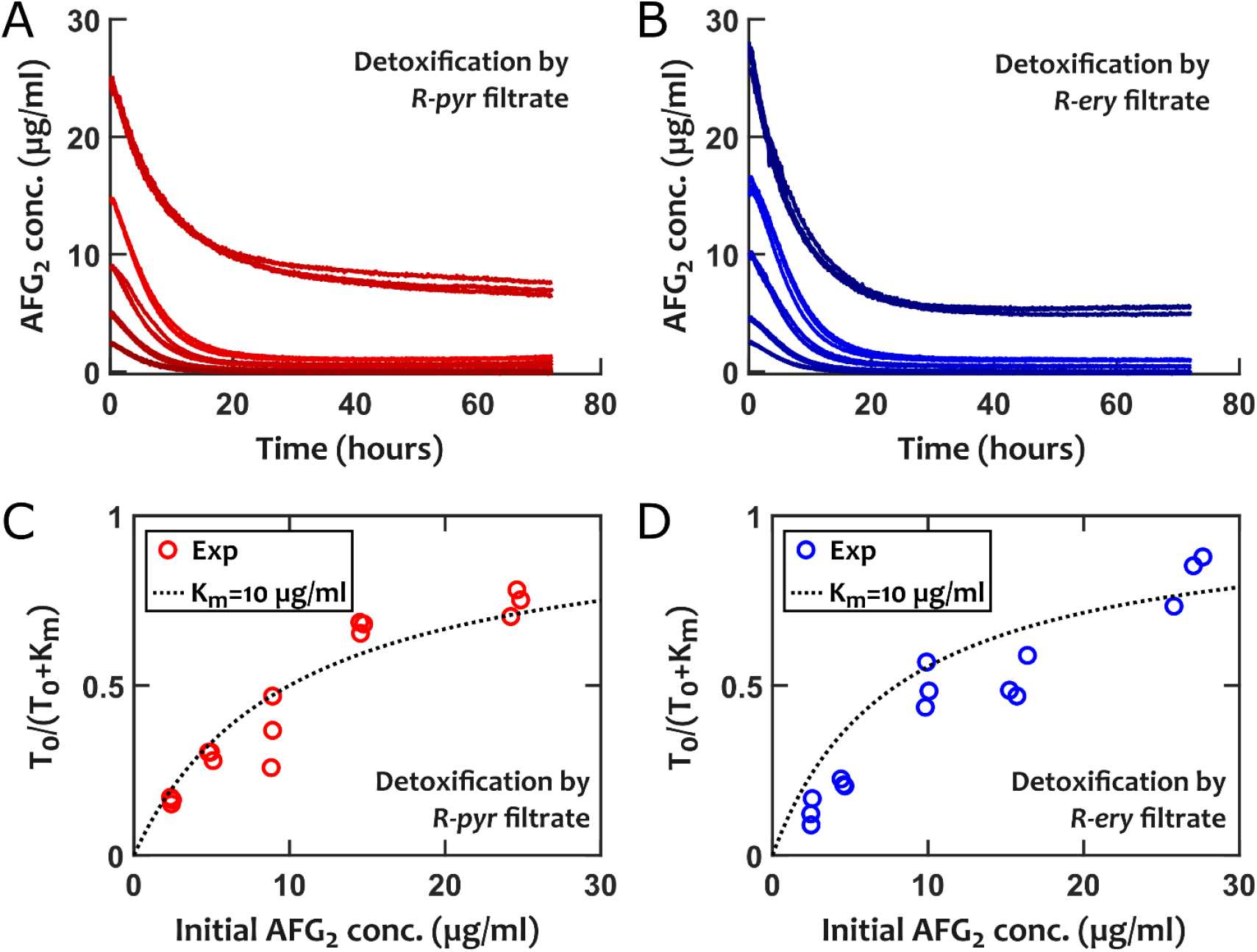
Monitoring detoxification rates, starting from different initial AFG_2_ concentrations allows us to estimate *K*_*m*_ for detoxification by *R-ery* and *R-pyr* filtrates. Detoxification kinetics of (A) *R-pyr* and (B) *R-ery* filtrates, starting from different initial concentrations of AFG_2_ are shown. (A) and (B) are used respectively to estimate ***T***_**0**_**/**(***T***_**0**_ + ***K***_***m***_) for detoxification by (C) *R-pyr* and (D) *R-ery* filtrates, allowing us to estimate the effective ***K***_***m***_ (see Eq 1). ***T***_**0**_ and ***dT*/*dt*** are estimated between hours 0.5 and 1.5 to avoid condensation artifact below 0.5 hours and enzyme defect effects in later times.

**Fig S4.**
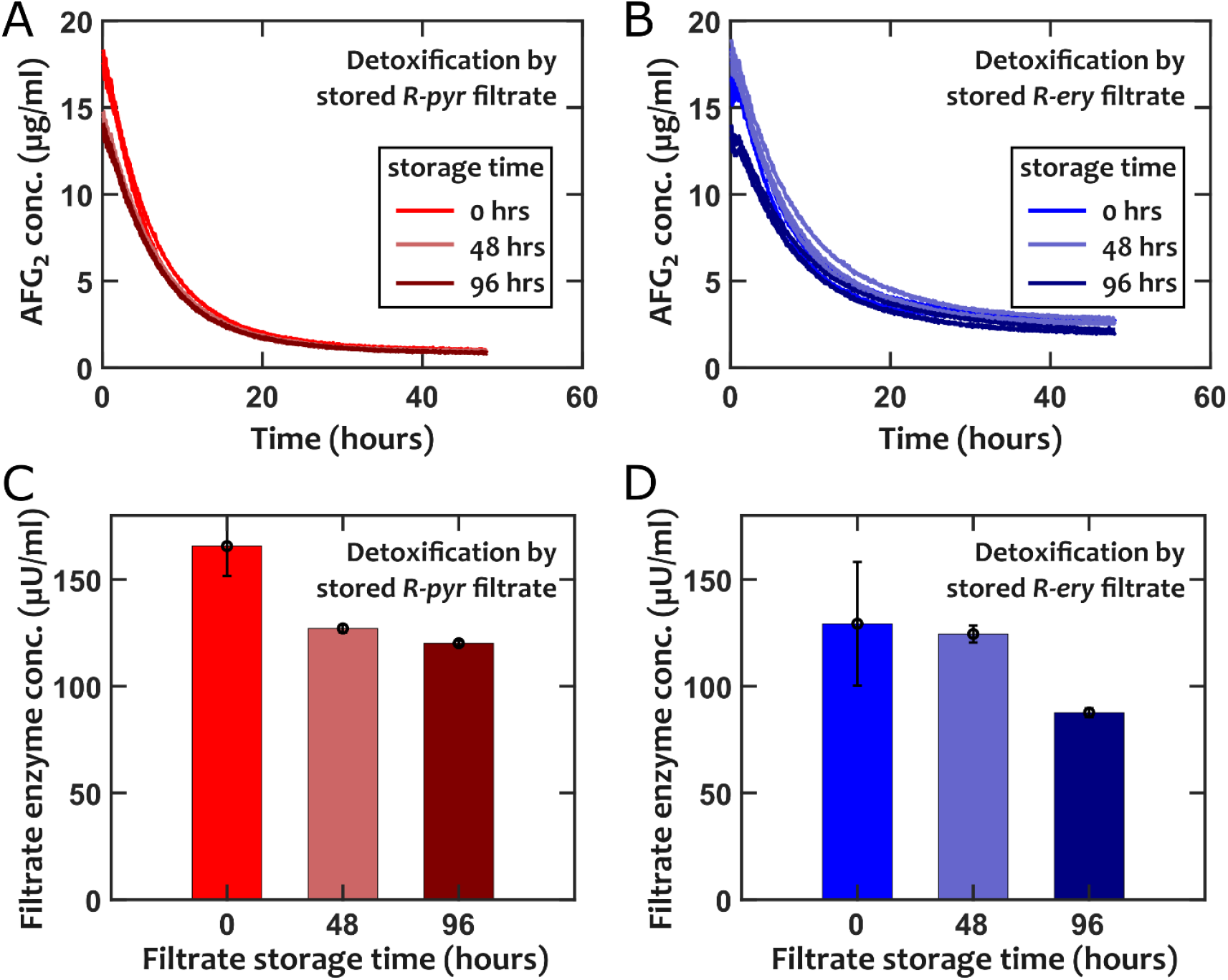
Comparing the detoxification of AFG_2_ by *R-ery* and *R-pyr* filtrates after storing them for different lengths of time shows only a modest drop in performance. Filtrates were stored in conditions similar to the detoxification assay (same treatment, 28°C, but no exposure to aflatoxin) for different durations (0, 48, or 96 hours) before using them for detoxification. The detoxification by (A) *R-pyr* and (B) *R-ery* stored filtrates were comparable to the original filtrate and enzyme decay during storage did not explain the significant slow-down in detoxification observed in our experiments (Fig 1). We also estimated the effective enzyme concentration in the stored filtrates (C-D), revealing only a modest decrease in detoxification after 96 hours of storage at 28°C. Error bars are s.d. among three replicates.

**Fig S5.**
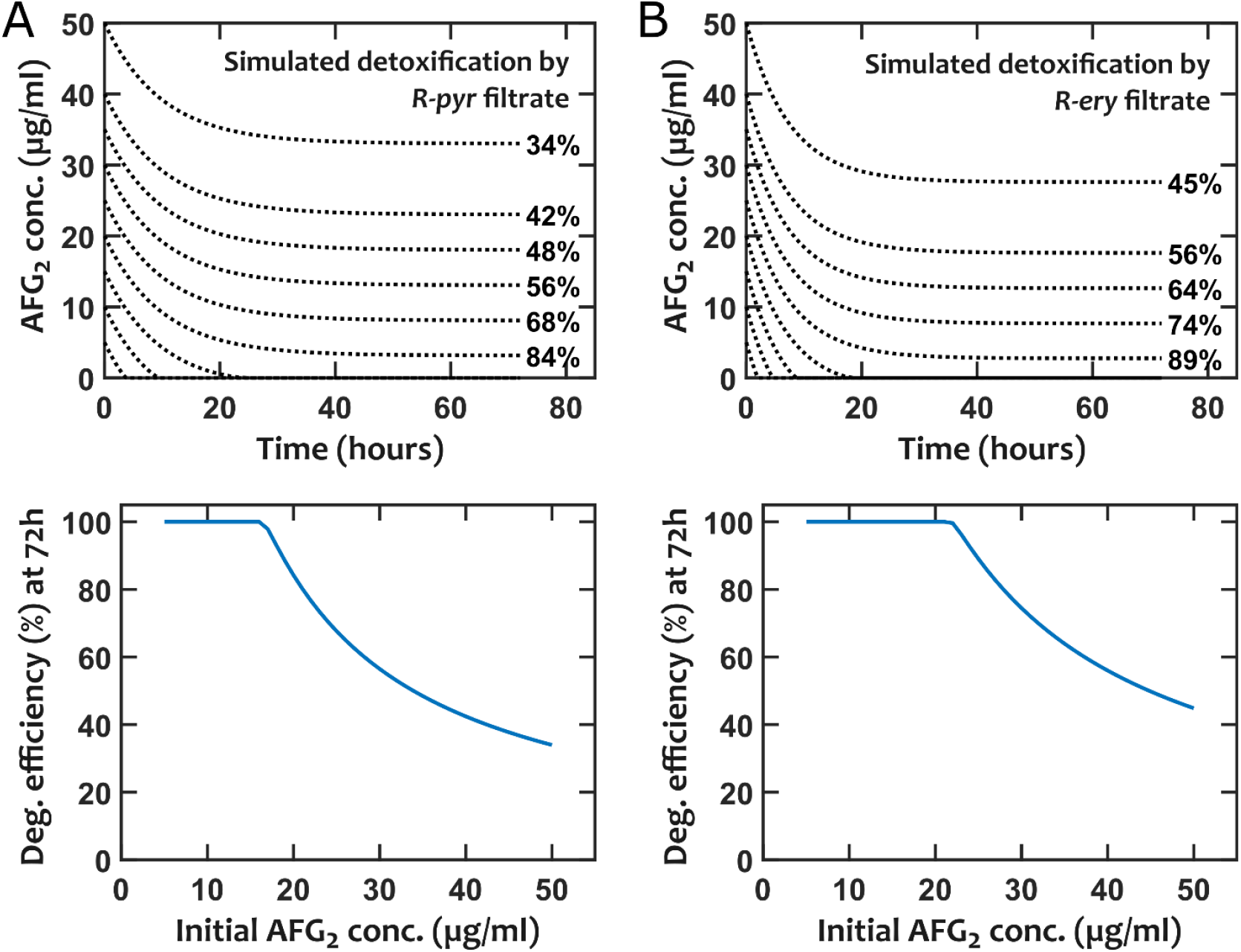
Simulated detoxification of AFG_2_ by *R-ery* and *R-pyr* filtrates shows that degradation efficiency is sensitive to the initial concentration of the toxin and thus not an ideal measure for comparing different strains. The formulation in Eq 1 is used for these simulations, with experimentally measured values of initial enzyme concentration and enzyme lifetime (Fig 2).

**Fig S6.**
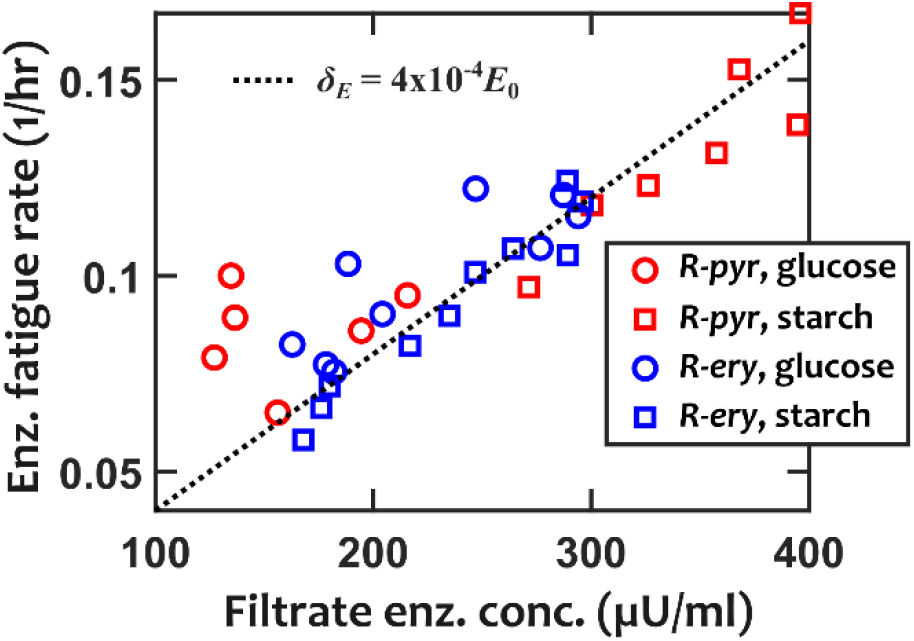
Enzyme fatigue rate (*δ*_*E*_) of *R-ery* and *R-pyr* filtrates show a consistent trend across species and environmental conditions (glucose versus starch). We estimate the relationship using a linear fit as *δ*_*E*_ = 1**/***τ* = 4 × 10^−4^*E*_0_.

**Fig S7.**
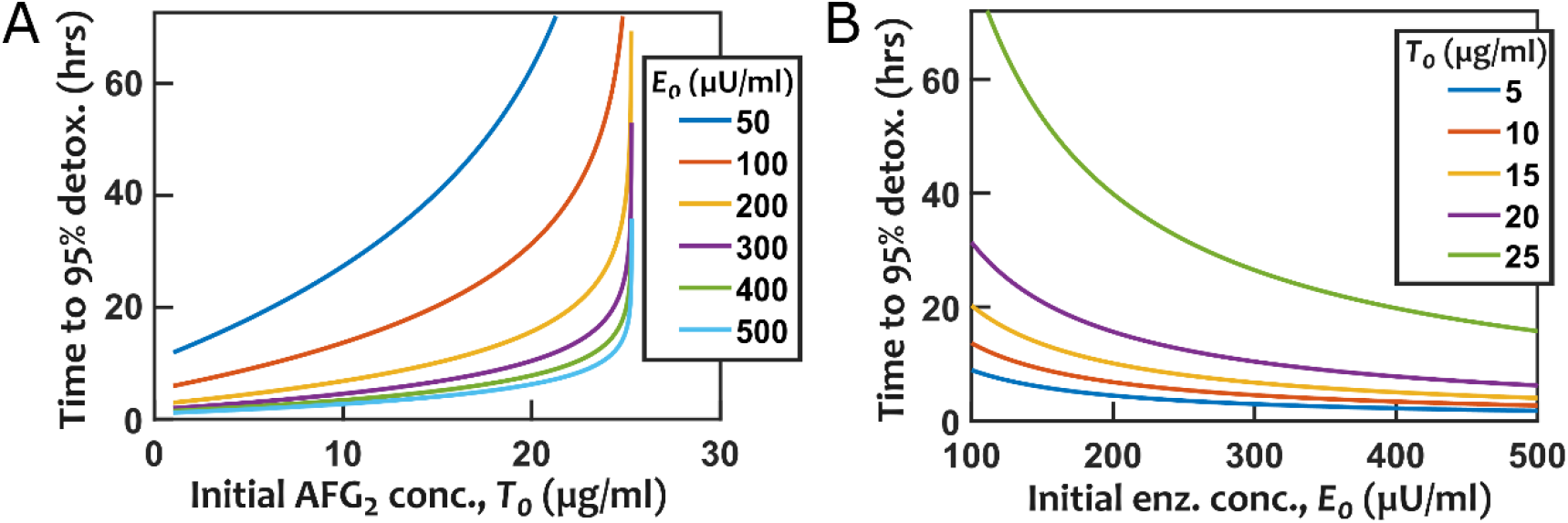
Simulated detoxification performance based on the time to detoxify 95% of the initial aflatoxin shows that enzyme fatigue limits the overall detoxification potential. (A) For each distinct enzyme concentration, detoxification time rapidly increases beyond a threshold aflatoxin concentration. (B) The detoxification of large aflatoxin concentrations exhibits diminishing returns at high enzyme concentrations. Eq (2) is used for these simulations, with the trade-off incorporated as a linear interpolation (Fig S6). These simulations are similar to Fig 6, except here K_m_ is assumed to be 3 µg/ml and *δ*_*E*_ = 1/*τ* = 6 × 10^−4^*E*_0_ to make the detoxification performance consistent with experimental data.

**Fig S8.**
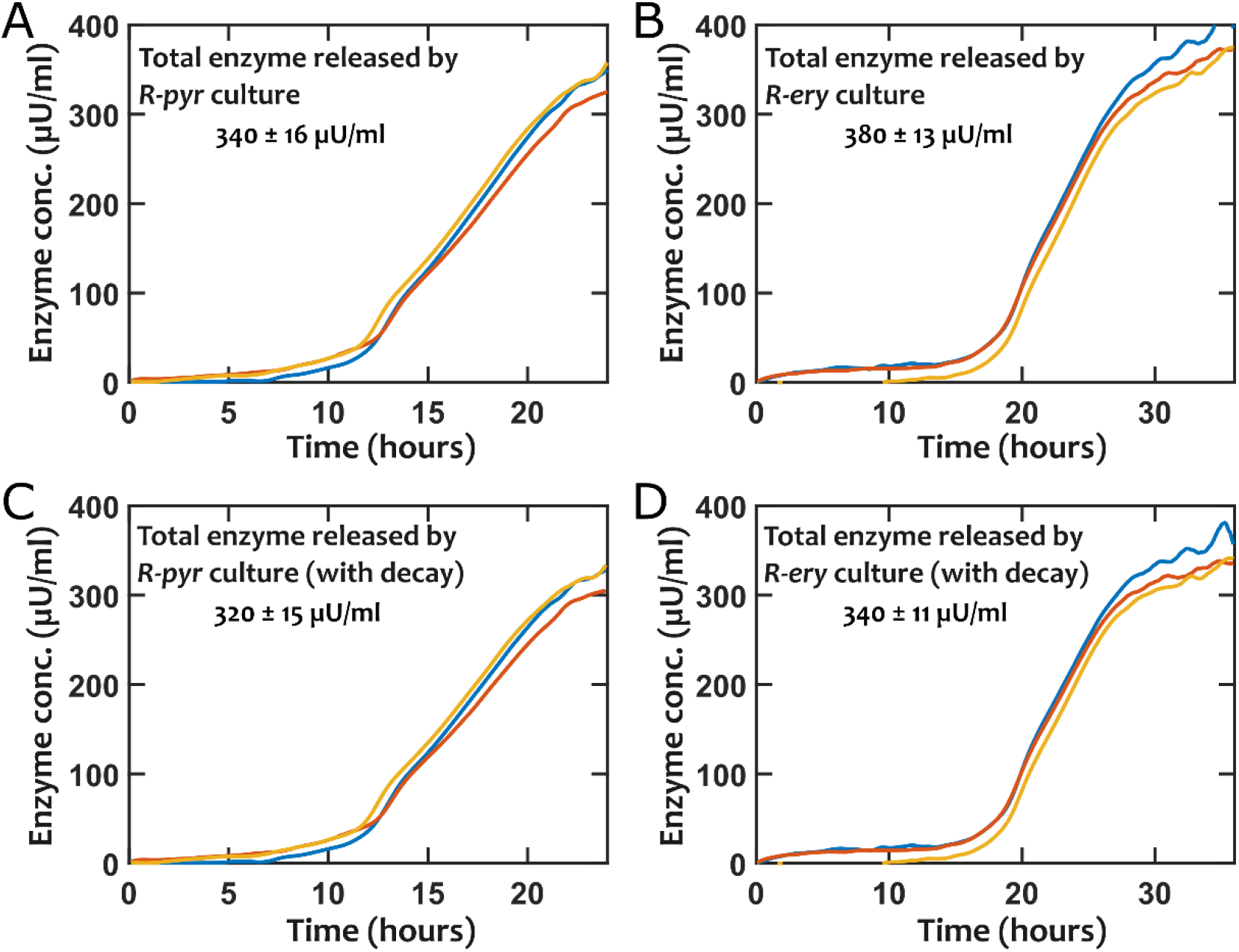
Estimated total released enzyme are inferred from live cultures of *R-pyr* and *R-ery* by integrating the enzyme released by cells (Figs 3 and 4) over time. Results for three replicates are shown. In (A) and (B) there is no enzyme decay. In (C) and (D), a decay rate of 0.008/hr is assumed, based on stored filtrate results in Fig S4. Overall, the estimated total released enzymes in live cultures in the presence of AFG_2_ were higher than those observed in filtrates obtained in the absence of AFG_2_ exposure (Fig 2).

## References

1. Bennett JW, Klich M. 2003. Mycotoxins. Clin Microbiol Rev 16:497–516.

2. Fernández-Cruz ML, Mansilla ML, Tadeo JL. 2010. Mycotoxins in fruits and their processed products: Analysis, occurrence and health implications. J Adv Res 1:113–122.

3. Kumar P, Mahato DK, Kamle M, Mohanta TK, Kang SG. 2017. Aflatoxins: A global concern for food safety, human health and their management. Front Microbiol 7:1–10.

4. Klingelhöfer D, Zhu Y, Braun M, Bendels MHK, Brüggmann D, Groneberg DA. 2018. Aflatoxin – Publication analysis of a global health threat. Food Control 89:280–290.

5. Amaike S, Keller NP. 2011. Aspergillus flavus. Annu Rev Phytopathol 49:107–133.

6. Wild CP, Gong YY. 2009. Mycotoxins and human disease: A largely ignored global health issue. Carcinogenesis 31:71–82.

7. Qian GS, Ross RK, Yu MC, Yuan JM, Gao YT, Henderson BE, Wogan GN, Groopman JD. 1994. A Follow-Up Study of Urinary Markers of Aflatoxin Exposure and Liver Cancer Risk in Shanghai, People’s Republic of China. Cancer Epidemiol Biomarkers Prev 3:3–10.

8. Wild CP, Montesano R. 2009. A model of interaction: Aflatoxins and hepatitis viruses in liver cancer aetiology and prevention. Cancer Lett 286:22–28.

9. Sarma UP, Bhetaria PJ, Devi P, Varma A. 2017. Aflatoxins: Implications on Health. Indian J Clin Biochem 32:124–133.

10. Womack ED, Brown AE, Sparks DL. 2014. A recent review of non-biological remediation of aflatoxin-contaminated cropsJournal of the Science of Food and Agriculture.

11. Adebo OA, Njobeh PB, Gbashi S, Nwinyi OC, Mavumengwana V. 2017. Review on microbial degradation of aflatoxins. Crit Rev Food Sci Nutr 57:3208–3217.

12. Vanhoutte I, Audenaert K, De Gelder L. 2016. Biodegradation of Mycotoxins: Tales from Known and Unexplored Worlds. Front Microbiol 7:561.

13. Sandlin N, Russell Kish D, Kim J, Zaccaria M, Momeni B. 2021. Computational biology for enzymatic removal of mycotoxins. OSF Prepr https://doi.org/10.31219/OSF.IO/9F4AN.

14. Udomkun P, Wiredu AN, Nagle M, Müller J, Vanlauwe B, Bandyopadhyay R. 2017. Innovative technologies to manage aflatoxins in foods and feeds and the profitability of application – A review. Food Control 76:127–138.

15. Shantha T. 1999. Fungal degradation of aflatoxin B1. Nat Toxins 7:175–178.

16. Samuel MS, Sivaramakrishna A, Mehta A. 2014. Degradation and detoxification of aflatoxin B1 by Pseudomonas putida. Int Biodeterior Biodegradation 86:202–209.

17. Zhao LH, Guan S, Gao X, Ma QG, Lei YP, Bai XM, Ji C. 2011. Preparation, purification and characteristics of an aflatoxin degradation enzyme from Myxococcus fulvus ANSM068. J Appl Microbiol 110:147–155.

18. Topcu A, Bulat T, Wishah R, Boyacı IH. 2010. Detoxification of aflatoxin B1 and patulin by Enterococcus faecium strains. Int J Food Microbiol 139:202–205.

19. Risa A, Krifaton C, Kukolya J, Kriszt B, Cserháti M, Táncsics A. 2018. Aflatoxin B1 and Zearalenone-Detoxifying Profile of Rhodococcus Type Strains. Curr Microbiol 75:907–917.

20. Cserháti M, Kriszt B, Krifaton C, Szoboszlay S, Háhn J, Tóth S, Nagy I, Kukolya J. 2013. Mycotoxin-degradation profile of Rhodococcus strains. Int J Food Microbiol 166:176–185.

21. Alberts JFF, Gelderblom WCACA, Botha A, van Zyl WHH. 2009. Degradation of aflatoxin B1 by fungal laccase enzymes. Int J Food Microbiol 135:47–52.

22. Risa A, Divinyi DM, Baka E, Krifaton C. 2017. Aflatoxin B1 detoxification by cell-free extracts of Rhodococcus strains. Acta Microbiol Immunol Hung 64:423–438.

23. Haskard CA, El-Nezami HS, Kankaanpää PE, Salminen S, Ahokas JT. 2001. Surface Binding of Aflatoxin B1 by Lactic Acid Bacteria. Appl Environ Microbiol 67:3086–3091.

24. Kim S, Lee H, Lee S, Lee J, Ha J, Choi Y, Yoon Y, Choi KH. 2017. Invited review: Microbe - mediated aflatoxin decontamination of dairy products and feeds. J Dairy Sci 100:871–880.

25. Martínková L, Uhnáková B, Pátek M, Nešvera J, Křen V. 2009. Biodegradation potential of the genus Rhodococcus. Environ Int 35:162–177.

26. Kuyukina MS, Ivshina IB. 2010. Application of Rhodococcus in Bioremediation of Contaminated Environments, p. 231–262. In. Springer, Berlin, Heidelberg.

27. Hashimoto Y, Nishiyama M, Yu F, Watanabe I, Horinouchi S, Beppu T. 1992. Development of a host-vector system in a Rhodococcus strain and its use for expression of the cloned nitrile hydratase gene cluster. J Gen Microbiol 138:1003–1010.

28. Sallam KI, Tamura N, Tamura T. 2007. A multipurpose transposon-based vector system mediates protein expression in Rhodococcus erythropolis. Gene 386:173–182.

29. Liang Y, Yu H. 2021. Genetic toolkits for engineering Rhodococcus species with versatile applications. Biotechnol Adv 49:107748.

30. Teniola ODD, Addo PAA, Brost IMM, Färber P, Jany K-DD, Alberts JFF, Van Zyl WHH, Steyn PSS, Holzapfel WHH. 2005. Degradation of aflatoxin B1 by cell-free extracts of Rhodococcus erythropolis and Mycobacterium fluoranthenivorans sp. nov. DSM44556T. Int J Food Microbiol 105:111–117.

31. Alberts J, Engelbrecht Y, Steyn P, Holzapfel W, Zyl, Van W. 2006. Biological degradation of aflatoxin B1 by Rhodococcus erythropolis cultures. Int J Food Microbiol 109:121–126.

32. Wu Q, Jezkova A, Yuan Z, Pavlikova L, Dohnal V, Kuca K. 2009. Biological degradation of aflatoxins. Drug Metab Rev 41:1–7.

33. Ellis WO, Smith JP, Simpson BK, Oldham JH, Scott PM. 1991. Aflatoxins in food: Occurrence, biosynthesis, effects on organisms, detection, and methods of control. Crit Rev Food Sci Nutr 30:403–439.

34. Zaccaria M, Dawson W, Kish DR, Reverberi M, Bonaccorsi Di Patti MC, Domin M, Cristiglio V, Dellafiora L, Gabel F, Nakajima T, Genovese L, Momeni B. 2021. Mechanistic Insight from Full Quantum Mechanical Modeling: Laccase as a Detoxifier of Aflatoxins. bioRxiv 2020.12.31.424992.

